# Multiplexed ^129^Xe HyperCEST MRI detection of genetically-reconstituted bacterial protein nanoparticles in human cancer cells

**DOI:** 10.1101/599118

**Authors:** Ryota Mizushima, Kanako Inoue, Hideaki Fujiwara, Atsuko H. Iwane, Tomonobu M. Watanabe, Atsuomi Kimura

## Abstract

Gas vesicle nanoparticles (GVs) are gas-containing protein assemblies expressed in bacteria and archaea. Recently, GVs have gained considerable attention for biotechnological applications as genetically-encodable contrast agents for MRI and ultrasonography. However, at present, the practical use of GVs is hampered by a lack of robust methodology for their induction into mammalian cells. Here, we demonstrate the genetic reconstitution of protein nanoparticles with characteristic bicone structures similar to natural GVs in a human breast cancer cell line KPL-4, and genetic control of their size and shape through expression of reduced sets of humanized gas vesicle genes cloned into Tol2 transposon vectors, referencing the natural gas vesicle gene clusters of the cyanobacteria *planktothrix rubescens/agardhii.* We then report the utility of these nanoparticles as multiplexed, sensitive and genetically-encoded contrast agents for hyperpolarized xenon chemical exchange saturation transfer (HyperCEST) MRI.

## Introduction

Gas vesicle nanoparticles (GVs) are gas-containing spindle- (or bicone-) shaped protein nanostructures with dimensions ranging from tens to hundreds of nm that are expressed in the cyanobacteria, algae and gram-positive bacteria. Since only gas molecules are permeable to the GV protein shell, GV-containing organisms acquire buoyancy by selective permeation of various ambient gases in the GV interior, which facilitates optimal supply of light and nutrition ^1-3^. While GVs have been studied in the field of microbiology for several decades, they have gained significant attention in recent years for their potential use for antigenic peptide display in vaccination ^4^, and as genetically-encodable contrast agents for ultrasonography ^5 6 7^ and HyperCEST MRI ^8,9^. Among these applications, HyperCEST MRI, which utilizes laser-polarized xenon-129 (HPXe) and chemical exchange saturation transfer (CEST) ^10^ to yield unprecedented enhanced MRI detection sensitivity, is of particular interest as GVs hold the potential to enable functional molecular imaging that is unfeasible in conventional thermally-polarized proton MRI. (In principle, GV concentrations of as low as pM - nM can be detected.) In this context, GVs can be thought of as an MRI analog of the green fluorescent protein (GFP) optical imaging reporter; with a genetically-encodable nature and multiplexing capability facilitated by ready modulation of their size and shape, similar to the multi-color variant of GFPs ^8^. However, despite their attractive features for imaging cellular / molecular processes, the practical *in-vivo* use of GVs as genetically-encoded contrast agents is at present hampered by a lack of robust techniques to introduce GVs into mammalian cells, which has been considered challenging due to the complexity of GV gene clusters ^11^.

GVs are composed of multiple proteins, and the number of genes responsible for GV expression is usually 8 – 14 (typically denoted GvpA, B, C etc.). Among these genes, the principal component proteins are the hydrophobic major protein GvpA and hydrophilic minor protein GvpC; the roles of other accessary GV genes in constituting GV wall structure remains a subject of controversy ^3^. In order to optimize GV delivery *in-vivo*, understanding of the minimal number and type of GV gene components required to reconstitute the GV nanostructure in mammalian cells is crucial. In this study, we focus on the unique gene clusters of GVs derived from *Planktothrix rubescens*/*agardhii* (praGV). The praGV gene clusters have been studied extensively by Walsby and coworkers ^12-14^, who showed that parts of praGV gene clusters are composed of *gvpA* and three variants of *gvpC* named *gvpC16*, *gvpC20* and *gvpC28*, though it is known that existence of other accessary GV genes have also been demonstrated ^3^. Properties of praGVs such as size, shape and stiffness (pressure threshold for collapse) are known to be modulated depending on the combination of *gvpC* variants included in their constituent gene clusters ^13^. Thus, we hypothesized that combinatorial expression of such reduced sets of genes in mammalian cells would allow reconstitution of protein nanoparticles with similar properties to GVs in natural organisms which can be functionalized as a contrast agent for HyperCEST MRI in mammalian cells and genetic control of their size and shape.

## Results and Discussion

In this work, we established stable expression of protein nanoparticles with characteristic bicone structures similar to natural GVs in human cancer cells and demonstrated genetic modulation of their size and shape. In addition, GVs were shown to be applicable as multiplexed, genetically-encoded HyperCEST MRI contrast agents in human cells *in-vitro*.

Firstly, praGV gene sequences available on the online database GenBank were examined (see **Supplementary Text**). The genes used in this study are listed in Table 1. These praGV genes with codons optimized for expression in mammalian hosts were synthesized, and cloned into Tol2 transposon vectors ^15,16 17^ under the control of tetracycline-inducible elements (Tet-On) ^18^. Puromycin-resistant genes were also included in the vector to select correctly transfected cells. T2A ^19^-fluorescent protein (EGFP, mKO2 ^20^ and mKate2 ^21^) fusion genes were cloned to the 3’ ends of GV genes (according to schemes in Figure 1a), for quantitative evaluation of GV gene expression by flow cytometry. T2A sequences were self-excised just after translation to avoid inhibiting GV structure formation, for example by steric hindrance of linked fluorescent proteins. We offer expression vectors of these humanized praGV genes at the National Bio-resource Center (https://dnaconda.riken.jp/search/depositor/dep103337.html).

**Table 1.** GV genes of *planktothrix rubescens*/*agardhii* used in this study. The nucleotide lengths (in units of base pairs, bp), the origin (strain) of the genes and their GenBank IDs, are shown.

| GV genes | Length (bp) | Strain | GenBank ID |
| --- | --- | --- | --- |
| *gvpA* | 219 | *P. rubescens*<br>BC-Pla9303 | AJ132357.1 |
| *gvpC16* | 420 | *P. rubescens*<br>BC-Pla9401 | AJ132354.1 |
| *gvpC20* | 519 | *P. rubescens*<br>BC-Pla9401 | AJ132361.1 |
| *gvpC28* | 738 | *P. agardhii*<br>CYA 29 | AJ253131.1 |

**Figure 1.**
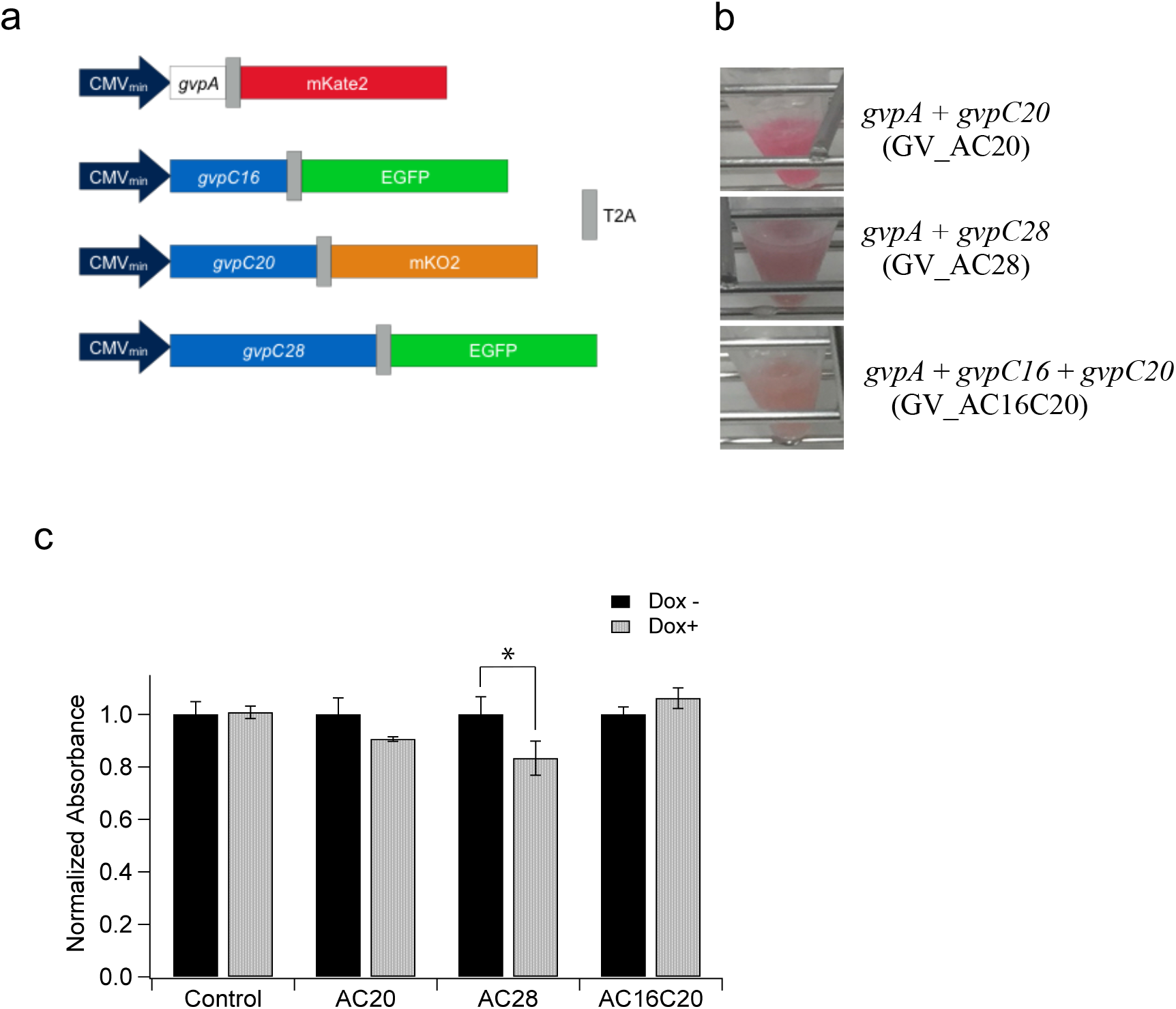
GV gene constructs, monoclonal KPL-4 cells expressing the GV gene sets and cell proliferation assay of the cells. (**a**) Schematics of the inserts of Tol2 vectors for transfection of humanized GV genes *gvpA*, *gvpC16*, *gvpC20* and *gvpC28*. CMV_min_: CMV minimal promoter. (**b**) Monoclonal KPL-4 cells stably expressing GV gene combinations in the presence of 1 µg/ml Dox, shown after pelleting. From top to bottom, monoclonal cells stably expressing: *gvpA* and *gvpC20* (GV_AC20), *gvpA* and *gvpC28* (GV_AC28), *gvpA*, *gvpC16* and *gvpC20* (GV_AC16C20). (**c**) Effect of GV gene expression on cell proliferation of each monoclonal cell, probed using a WST-8 reagent at 72 hours after addition of Dox (Dox+) or an equivalent amount of PBS (Dox-) into the culture medium. Normalized absorbances of the average of 8 separate cultures of each monoclonal cell are shown. An asterisk (*) indicates statistical significance: *p* < 0.05, t-test. Error bars indicate S.E.M..

Combinations of praGV gene constructs were co-transfected into KPL-4 human breast cancer cells ^22^. Combinations were chosen in reference to the natural *Planktothrix rubescens*/*agardhii* GV gene clusters ^13^. In this study, we investigated 3 specific combinations: *gvpA* + *gvpC20* (GV_AC20), *gvpA* + *gvpC28* (GV_AC28) and *gvpA* + *gvpC16* + *gvpC20* (GV_AC16C20). Transfected cells expressing the GV gene sets at high levels were selected using puromycin selection and fluorescence-activated cell sorting (FACS). Monoclonal single colonies were picked and expanded for GV_AC20, GV_AC28 and GV_AC16C20 cells. Obtained monoclonal cell pellets are shown in Figure 1b. Expression levels of the fluorescent proteins of the monoclonal cultures were measured by flow cytometry (Supplementary Figure 1). The expected cellular burden to express these GV genes in mammalian cells was evaluated by a cell proliferation assay using a WST-8 reagent (**Materials and Methods**). GV_AC28 cells showed a significant decrease of cell proliferation after 72 hours of induced GV gene expression with doxycycline (Dox+) (*p* = 0.024, t-test) compared with that of non-induced control cells (Dox−) (Figure 1c). Other GV cells did not show a significant decrease of cell proliferation (*p* = 0.86, 0.18 and 0.34, t-test for control, GV_AC20 and GV_AC16C20 cells, respectively).

To confirm expression of foreign protein nanoparticles in the cells, we developed a method of purifying buoyant protein nanoparticles from mammalian cells (Figure 2a). The protocol was modified from that previously described ^23^ to exploit the much lower density of GVs compared to water. The purified putative nanoparticle suspensions were observed by TEM. Characteristic bicone structures were observed when using a standard negative staining protocol (**Materials and Methods**) in a purified suspension of the GV_AC28 cells (Figure 2b). In other cell clusters, distinct structures were not observed. Thus, we tested another protocol, usually adopted for observation of intracellular structures in cells, which we proposed would help avoid collapse of the nanoparticles during TEM observation. Putative nanoparticle suspensions were first embedded in ~ 1 mm^3^ 2% agarose gels and fixed with glutaraldehyde (GA) (**Materials and Methods**). Using this protocol, distinct bicone structures were also observed in suspensions purified from GV_AC20 and GV_AC16C20 cells (Figure 2b). It is noteworthy that these observed nanostructure were not cellular organelles or any membranous structures, because they were collapsed by the act of osmotic pressure and detergent used in the purification method. We additionally note that the TEM images could not be any salt crystals because applied staining method specifically contrasted the proteins. We also obtained Dynamic Light Scattering data on the purified nanoparticle suspensions to gain insights into the size distribution of the protein nanoparticles (Supplementary Figure S2). The obtained TEM images and DLS data clearly indicated the expression of characteristic bicone nanostructures in mammalian cells for each of our GV gene constructs, and genetic control of their size and shape was observed, as hypothesized (**Supplementary Text**). In addition, the cell proliferation assay indicated that cellular burden of these nanoparticle expression in mammalian cells may depend on the expressed nanoparticle size (Figure 1c).

**Figure 2.**
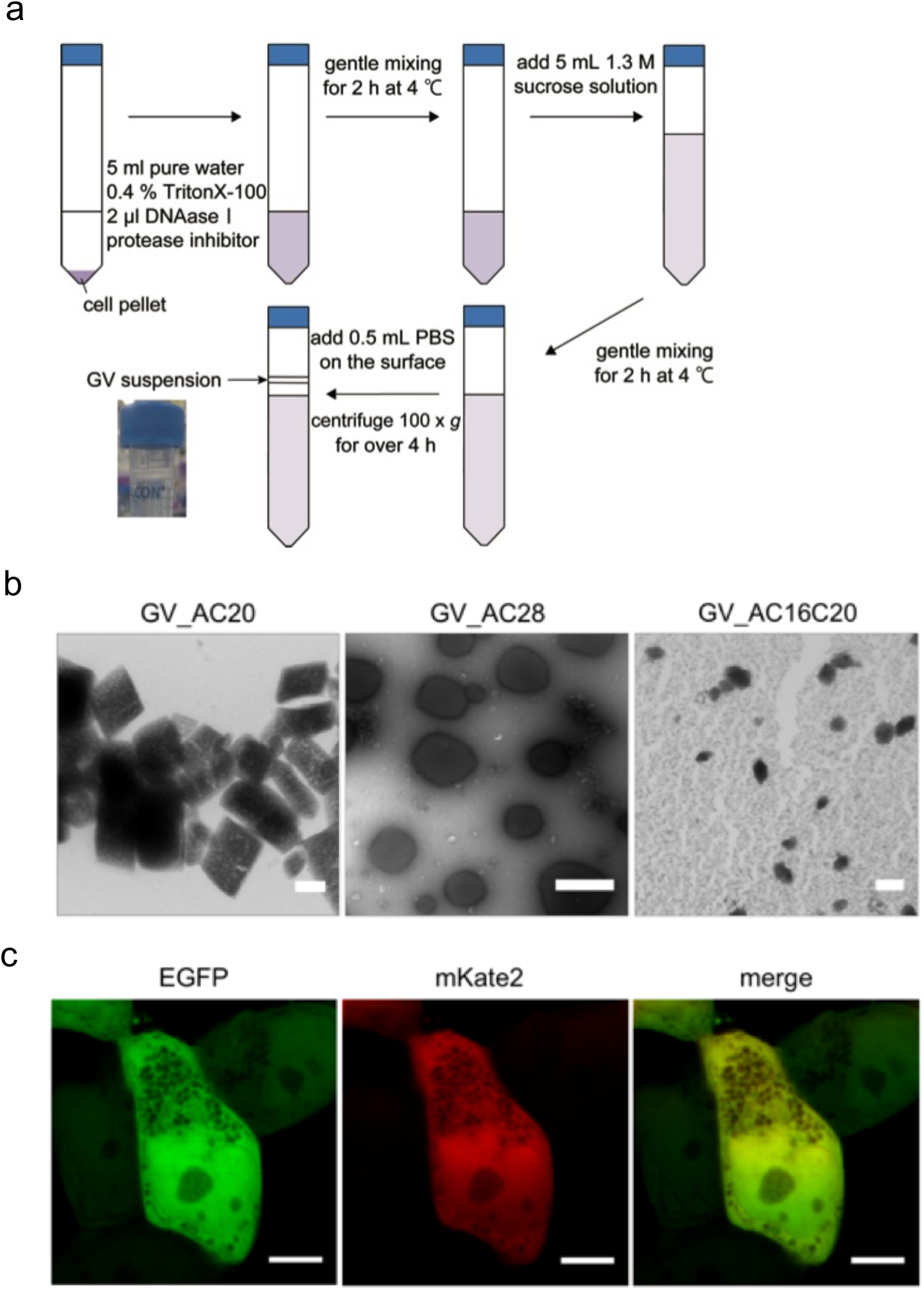
Expression of GVLPs in human cells. (**a**) Schematic illustration of the protocol developed to purify GVLPs from mammalian cells. (**b)** TEM images of purified GVLPs from the cells: GV_AC20 (left), GV_AC28 (middle) and GV_AC16C20 (right). Scale bars indicate 100 nm (left), 500 nm (middle) and 100 nm (right). (**c**) Confocal microscopy images of GV_AC28 cells. Fluorescence images of EGFP (left), mKate2 (middle) and merged image of the two (right). Scale bars indicate 5 μm in all three images.

Moreover, we were able to clearly observe the nanostructures in GV_AC28 cells as a negative contrast by fluorescent confocal microscopy due to their large size (above the spatial resolution of confocal microscopy) (Figure 2c). We call these nanoparticles expressed in mammalian cells by reduced humanized GV gene sets with characteristic bicone nanostructures similar to natural GVs GV-like particles (GVLPs) and our proposed methodology of GVLPs expression in mammalian cells with genetic control of their size and shape utilizing humanized praGV genes as FC-SOPRAGA: <u>F</u>acilitated <u>C</u>ontrol of <u>S</u>elf-<u>O</u>rganized <u>P</u>lanktothrix <u>R</u>ubescens/<u>A</u>gardhii <u>G</u>as vesicle -like particle <u>A</u>ssembly.

To investigate whether the reconstituted GVLPs could be used as multiplexed and genetically-encoded HyperCEST MRI contrast agents (as previously demonstrated in prokaryotic cells ^8^), we measured the MR signal intensities of hyperpolarized xenon dissolved in cells expressing GVLPs and control KPL-4 cells as a function of saturation frequency offset (i.e. recorded CEST Z-spectra) using a 2 s saturation time. The saturation was performed to detect the chemical shift of GVLP-bound xenon, i.e. the saturation chemical shift at which the bulk dissolved xenon signal was most attenuated by chemical exchange transfer of saturated spins between GVLPs and bulk dissolved xenon.

The HyperCEST Z-spectrum of GV_AC28 cell samples consistently showed a maximal signal decrease at around 20 ppm saturation offset chemical shift, and occasionally minor saturation peaks at around 70 ppm, in addition to the bulk dissolved phase HPXe saturation peak at 194 ppm, while the control KPL-4 cells showed only the 194 ppm saturation peak of bulk dissolved xenon with a broad and weak background around 0 ppm (Figure 3a). We believe the two unique peaks of GVLP-bound xenon may be explained by the inhomogeneity of GVLP size in GV_AC28 cells as indicated in TEM images (Figure 2b) and DLS data (Supplementary Figure 2). The 20 ppm peak may be attributed to the xenon bound to large vesicles and the minor peak (~70 ppm) presumably relates to the xenon bound to smaller vesicles ^24^. Furthermore, we observed a slight, but repeatable saturation peak of GV_AC16C20 cells at around 60 ppm in its Z-spectrum despite a lower number of GV cells (2.0 × 10^7^ cells / ml vs. 8.0 × 10^7^ cells / ml for GV_AC28 cells). In this case, the background signal in the Z-spectrum was reduced, similar to that previously reported for the β-lactamase HyperCEST reporter gene ^11^. Therefore, we concluded that the peak at 60 ppm was indeed derived from the saturated GV-bound xenon transfer rather than an experimental artifact (Figure 3a). On the other hand, the Z-spectrum of GV_AC20 cells (Supplementary Figure 3) did not show any practical saturation peaks for HyperCEST contrast at 2 s saturation time.

**Figure 3.**
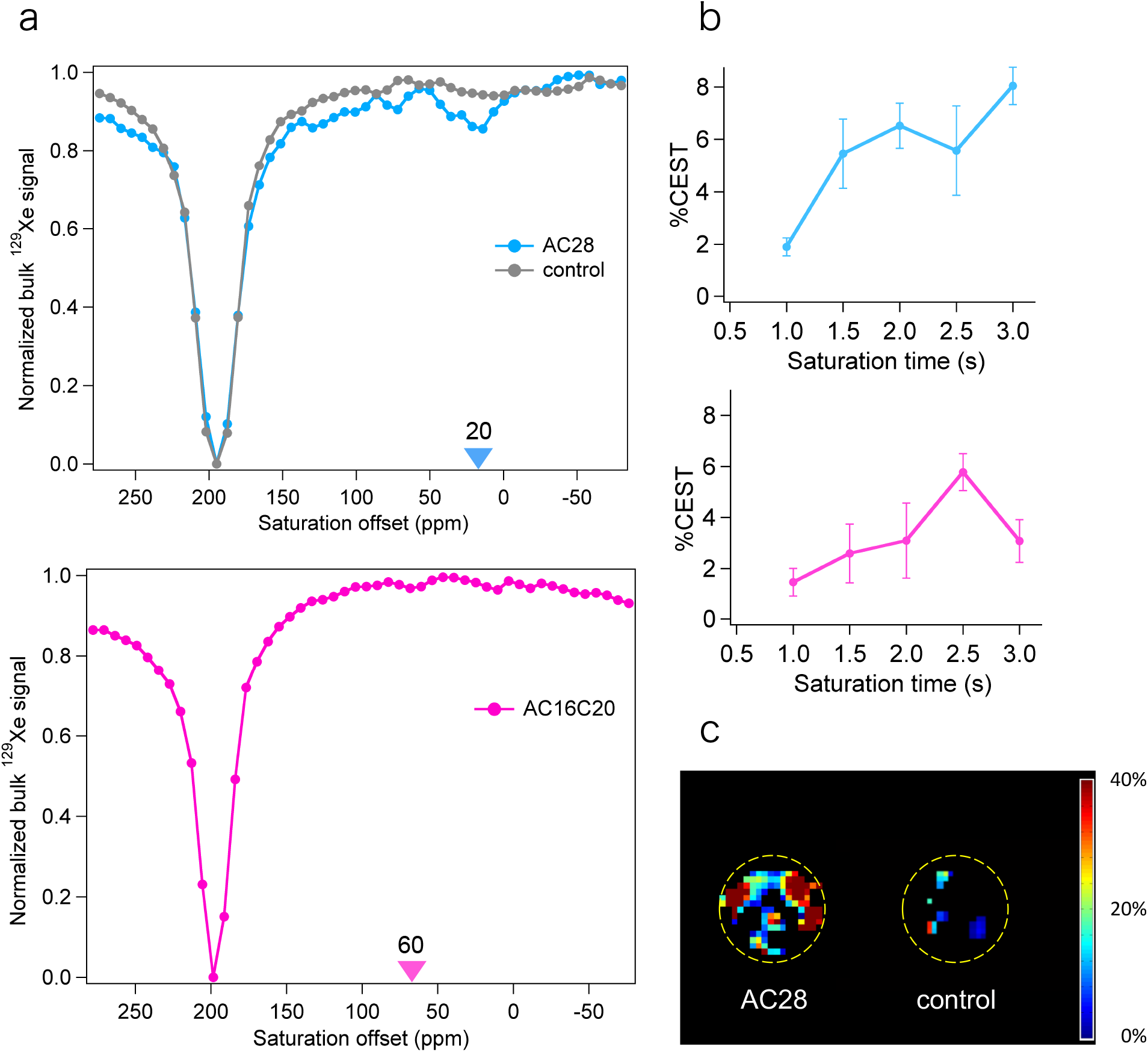
HyperCEST MR Z-spectra and images of GV cells. The strengths of saturation RF pulses employed for the HyperCEST MR Z-spectra and images were 1.5 μT (30dB) and 5.9 μT (52dB), respectively. (**a**) Z-spectrum of GV_AC28 cell and control KPL-4 cell samples with 8.0 × 10^7^ cells / ml (upper) and GV_AC16C20 cell sample with 2.0 × 10^7^ cells / ml (lower). Color triangles indicate the saturation offsets used to calculate %CEST contrast in (b). (**b**) Saturation time dependency of %CEST contrast for GV_AC28 cells (upper) and GV_AC16C20 cells (lower). (**c**) HyperCEST MR images of GV_AC28 cell and control KPL-4 cell samples with 8.0 × 10^7^ cells / ml. The dotted yellow line indicates the wall of the 10φ glass tube in which the cell suspension was contained. The color bar indicates %CEST contrast.

Subsequently, bulk dissolved xenon NMR signal intensity was measured after applying saturation pre-pulses at the on-resonance and off-resonance chemical shifts (on-resonance ^signal intensity *Son* and off-resonance signal intensity *Soff*, respectively) as a function of^ saturation time and the %CEST contrast was calculated as defined in the equation for GV_AC28 and GV_AC16C20 cells (Figure 3b):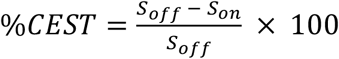.

In GV_AC28 cells, chemical shifts of on-resonance and off-resonance saturation pre-pulses were set to 20 ppm and −20 ppm, respectively, and for GV_AC16C20, 60 ppm and −60 ppm were used. GV_AC28 cells showed 5.5 ± 1.3 and 6.5 ± 0.9 % CEST contrast at 1.5 s and 2 s saturation time, respectively. At 3 s saturation time, CEST contrast maximally reached 8.0 ± 0.7 %. For GV_AC16C20 cells, 2.6 ± 1.1 and 3.1 ± 1.5 % CEST contrast was observed at 1.5 s and 2 s saturation time, with a maximal 5.8 ± 0.7 % contrast at 2.5 s saturation time. Considering the longitudinal relaxation time (T^1^) of gaseous xenon *in vivo* (~ 30 s, note that the T^1^ of dissolved xenon is lower), the saturation time for in-vivo HyperCEST MRI should ideally be less than 2 s ^25,26^. It is noteworthy that a low saturation time is also desirable for decreased specific absorption rate, which is an important consideration regarding the application of GVs in human patients. Taking these concerns into account, a HyperCEST MR image of GV_AC28 cells was successfully acquired by applying a 2 s saturation scheme at 20 ppm (Figure 3c).

## Conclusions

In this study, we showed that expression of reduced GV gene sets derived from *planktothrix rubescenes/agardhii* in mammalian cells resulted in the formation of GVLPs in the cell and they could be functionalized as a genetically-encoded HyperCEST MRI contrast agents. GVLPs have distinct properties from that of natural GVs expressed in bacterial cells and it should be further characterized in the future. A recent study^27^ also demonstrated that gas vesicles can be expressed in mammalian cells and their utility as a genetically-encoded ultrasound contrast agent by similar methods under almost the same overall experimental design, which is consistent with our results. Note that our method of mammalian expression of gas vesicles was conceived independently of the previous study^27^ demonstrated by the patent application already open in public^28^. GV_AC28 construct demonstrated promising CEST-based MRI contrast within a practical saturation time, which has not been reported to date in genetically-encoded HyperCEST MRI contrast agents for mammalian cells. Our findings should help facilitate the future development of GV-based HyperCEST MRI reporters for *in-vivo* applications, though realization of *in-vivo* HyperCEST still requires overcoming several technical hurdles, such as continuous or high-volume batch supply of xenon with non-renewable polarization^29^ and tissue-specific targeted delivery of GVs.

## Acknowledgments

We thank Dr. Akira Takai and Prof. Yasushi Okada of RIKEN for donating Tol2 transposon vectors and for providing advice regarding their use. We thank Prof. Kaoru Mitsuoka of Osaka University and Ms. Rina Nagai of RIKEN for their support of TEM observations of GVs. We thank Prof. Takashi Jin of RIKEN for his kind offer of DLS machine time. We thank Dr. Kazuko Okamoto and Ms. Keiko Yoshizawa of RIKEN for their technical assistance with FACS operation, cell culture and molecular cloning. We also thank Dr. Neil J Stewart for proofreading the manuscript.

We acknowledge Kawasaki Medical School and Prof. Junichi Kurebayashi for generously providing KPL-4 cells. This work was supported by the NIMS microstructural characterization platform, a program of the "Nanotechnology Platform" of the Ministry of Education, Culture, Sports, Science and Technology (MEXT), Japan.

This work was supported in part by JSPS KAKENHI Grant Number JP17K20121.

## Author contributions

T.M.W. suggested project of GV expression in mammalian cells. R.M. conceived FC-SOPRAGA methodology. R.M. designed the GV gene constructs, cultured the cells, transfected vectors, produced GV cells and purified GVLPs from the cells. R. M. carried out cell proliferation assays and analyzed the data. R.M., K.I. and A.H.I. prepared the TEM samples and acquired TEM images of GVLPs. R.M. acquired FACS, confocal microscopy and DLS data sets. R.M. H.F. and A.K. designed the HyperCEST NMR/MRI experiments. A.K. acquired the HyperCEST NMR/MRI data sets. R.M and A.K. analyzed and interpreted the data. A.K. produced HyperCEST MR images. R.M. and A.K. wrote the manuscript.

## Competing interests

RIKEN has a pending patent application already open in public (WO2018043716A1) regarding this work, in which R.M. and T.M.W. are inventors.

## Data and materials availability

All data and materials underlying this study are available. Expression vectors of humanized praGV genes were deposited to and are available from the BioResource Research Center, RIKEN. The article was previously posted on bioRxiv (http://biorxiv.org/cgi/content/short/599118v1).

## MATERIALS AND METHODS

### Synthesis of humanized GV genes

GV genes were searched in Genbank over as many strains as possible derived from *planktothrix rubescens*/*agardhii* to synthesize humanized genes for mammalian expression of GV proteins. The gene *gvpA* was chosen from the strain pla-9303, *gvpC16* from pla-9401, *gvpC20* from pla-9401 of *Planktothrix rubescens* and *gvpC28* from *Planktothrix agardhii* CYA29. Coding sequences of these genes with codons optimized for expression in mammalian hosts were synthesized (outsourced to Genscript).

### Molecular cloning

Primers used for gene cloning were purchased from Hokkaido System Science. Coding sequences of GV genes were PCR-amplified with 5’ primers encoding a NheI site and 3’ primers encoding an EcoRI site without termination codons using KOD-plus-Neo (TOYOBO). T2A - fluorescent protein (mKate2, mKO2 and EGFP) fusion genes were also PCR amplified with 5’ primers encoding an EcoRI site and 3’ primers encoding a NotI site. A Tol2 cloning vector (donated by Dr. Akira Takai of RIKEN and described in detail previously ^16-18^) was also digested in the same way. The PCR products and restriction enzyme digestions were purified by agarose gel electrophoresis followed by processing with the Wizard SV Gel and PCR clean up system (Promega). Restriction enzymes were purchased from Fermentas. The digested PCR products of GV genes and T2A-fluorescent protein fusion genes were ligated to the vectors using Ligation high Ver.2 ligase (TOYOBO) following the recommended protocol of the manufacturer. Plasmids were prepared from the bacterial liquid culture by using the Pure Yield Plasmid Miniprep System (Promega) and PureLink HiPure Plasmid Midiprep Kit (Invitrogen). The DNA sequences were read by dye terminator cycle sequencing using the BigDye Terminator v3.1 Cycle Sequencing Kit (Applied Biosystems).

### Transfection and cell culture

Plasmid DNAs containing Tol2 vectors comprising GV genes and fluorescent protein genes described above and transposase mRNA were introduced to KPL-4 cells by lipofection using FuGene HD (Promega) in OptiMEM (Gibco), according to manufacturer protocols. Transfected cells were maintained in a low-glucose DMEM (FUJIFILM Wako Pure Chemical Corporation) with 10% FBS (Gibco) and 1% penicillin-streptomysine (Gibco).

### KPL-4 monoclonal cell lines expressing GV gene sets

KPL-4 cells were transfected with the GV gene constructs and 1 µg/ml puromycin (Invitrogen) was added to cell cultures to select only the transfected cells. The cells were sorted several times to collect polyclonal cells expressing GV gene sets at high levels (> × 10^2^ fluorescence intensity compared to control cells) by detecting fluorescence of mKate2, mKO2 and EGFP in correlation with the expression levels of GV genes using a BD FACS Aria Ⅲ cell sorter (BD Bioscience). Spectral overlaps of each fluorescence were calibrated using the cells expressing each single fluorescent protein gene. Among the polyclonal cells with high gene expression level, single colonies were picked and expanded to establish the monoclonal GV cell lines expressing the chosen GV gene set at a high level. Expression levels of each monoclonal cell line were checked by fluorescence intensity profiles (Supplementary Figure 1).

### Cell proliferation assay

Cell proliferations of each monoclonal GV cells and control KPL-4 cells with or without 1 μg/ml Dox were probed using a Cell Counting Kit-8 (WST-8 reagent) (Dojindo Molecular Technologies) ^28^. For this reagent, relative absorbance at 450 nm is proportional to the number of viable cells. ~ 5000 cells in 100 μl medium described above were prepared by a Countess Ⅱ cell counter (Invitrogen) and cultured in each well of a 96-well plate (IWAKI) with 10 μl addition of 10 μg/ml Dox or PBS for each GV cell culture and control KPL-4 cell culture. After 72 hours of incubation, 10 μl WST-8 reagent was added to each well and cultures were further incubated for an hour. Then, absorbance of each well at 450 nm was measured by a microplate reader (Thermo Fischer Scientific). 8 wells were used for each cell culture condition. Means and S.E.M.s of the absorbances of the 8 wells for each condition were calculated. Absorbance was normalized to that of the mean of the “Dox-” condition data for each GV cell culture and control cell culture.

### Purification of GVLPs from mammalian cells

Each monoclonal GV cell line was cultured in a 15 cm tissue culture dish (IWAKI) with 30 ml DMEM until it reached 100 % confluent growth. Cells were washed with PBS, peeled with 0.25 % trypsin-EDTA (Sigma-Aldrich) and collected in 50 ml sample tubes. Cells were centrifuged, collected and re-suspended in PBS. The samples were transferred to a new 15 ml sample tube, centrifuged again and cell pellets were collected. The cell pellets were suspended in 5 ml pure water at 4 ℃ with 0.4 % TritonX-100 (Sigma-Aldrich), 2 μl DNAase Ⅰ(Sigma Aldrich) and a protease inhibitor cocktail (Roche). The samples were incubated at 4 ℃ with gentle mixing for two hours. The same amount of 1.3 M sucrose solution (5 ml) was added to the sample (thus the final sucrose concentration was 0.65 M), which was then incubated for a further two hours with gentle mixing at 4 ℃. 0.5 ml PBS was then carefully placed on the surface of the sample. The samples were then centrifuged for over 4 hours at 100 × *g* at 4 ℃ with a swing bucket rotary and appropriate balance to keep the rotating samples perpendicular to the gravity axis. After the centrifugation, ~ 200 µl samples were collected from the sample surface and the GVLP suspension was used for subsequent analysis.

### Transmission Electron Microscopy (TEM)

Purified GVLP samples were prepared for TEM analysis in two ways: (1) Purified GVLP from GV_AC28 cells was fixed with the same amount of 2.5 % glutaraldehyde (GA) solution (FUJIFILM Wako Pure Chemical Corporation) and prepared with a standard negative staining protocol as follows. Purified suspension of 3 µl was placed onto the 200 mesh copper grid and incubated for 60 seconds. The extra solution was absorbed with filter paper and a water droplet of 3 µl was placed on the grid. The droplet was absorbed again immediately with filter paper, a 2 % uranium acetate droplet of 3 µl was placed on the grid and absorbed 3 times. The dried sample was then used for TEM JEM1011 (JEOL) at an acceleration voltage of 100 kV. (2) GV_AC20 cells and purified GVLPs from GV_AC20 cells and GV_AC16C20 cells were first embedded in 2 % agarose gels (~ 1 mm^3^) prior to fixation. After the fixation with half Karnovsky mixture (2.5 % GA and 2 % paraformaldehyde), the samples were dehydrated by soaking them with 30 %, 50 %, 70 %, 80 % and 90 % sequences for 10 minutes each. After that, samples were soaked three times with 100 % ethanol. Samples were then soaked with 100 % propylene oxide (PO) for 10 minutes and in 1:1 and 1:2 mixture of PO solvent and Quetol-812 (Nisshin EM) epoxy resin (EPON) for 6 hours and 2 hours, respectively. Furthermore, the samples were soaked in a mixture of EPON and epoxy curing agent DMP-30 for 6 hours. Finally, the samples were embedded in the same EPON and DMP-30 mixture formed with a plastic or silicon mold and cured in the oven at 45 ℃ for 24 hours and at 60 ℃ for 72 hours, respectively. The obtained resin block was trimmed with a glass knife and ultra-thin sections (~ 70 nm) were prepared using a Leica EM UC7 ultramicrotome (Leica) and ultra diamond knife 35°(Diatome). The resulting ultra-thin sections were scooped onto 200 mesh copper grids and stained with modified Sato’s lead solution and 4 % uranium acetate solution. The grids were dried and observed by TEM Hitachi-7500 (Hitachi High-Technologies) at an acceleration voltage of 120 kV or TEM JEM1011 (JEOL) at an acceleration voltage of 100 kV.

### Dynamic Light Scattering (DLS)

Size distributions of nanoparticles in the GVLP suspensions were measured at 25 ℃ by a Zetasizer Nano-ZS (Malvern Instruments) with a 633 He/Ne Laser. The distribution of hydrodynamic size of purified GVLPs was estimated from the DLS data (Supplementary Figure 2).

### Confocal Microscopy

The GV_AC28 cells incubated with Dox for over 3 days were plated onto a 35 mm glass bottom dish filled with phenol red-free DMEM (Gibco) with supplements. Fluorescence images were obtained by an inverted confocal microscope FV-1000 (Olympus) combined with a stage-top ^incubator INUC-KRi (Tokai-Hit) to control the temperature at 37 ℃. CO2 was loaded at 5 %^ into the incubator GM-2000 (Tokai-Hit). GFP and mKate2 were simultaneously excited by 473 nm and 559 nm lasers, respectively. A 60x objective lens (NA 1.40, oil) PLAPON 60XOSC2 (Olympus) and an observation area of 30.705 × 30.705 μm^2^ (2048 × 2048 pixels) were used.

### MR Measurements

MR measurements were performed on an Agilent Unity INOVA 400WB high-resolution NMR system (Agilent Technologies). A 9.4 T vertical magnet with a bore width of 89 mm (Oxford Instruments) was used. A multinuclear (^1^H-^129^Xe) NMR Probe (inner φ:10 mm) was used for spectroscopy and a self-shielded gradient probe Clear Bore DSI-1117 (inner φ: 34 mm) (Doty Scientific) for imaging. Litz volume RF coils tuned to the Larmor frequency of ^129^Xe (110.6 MHz) were employed for ^129^Xe detection. A gas mixture of 70% xenon (natural abundance, ^129^Xe fraction 0.26) and 30% N_2_ was hyperpolarized using a home-built continuous-flow type ^129^Xe polarizer, and delivered to the GV cell suspensions in a 10 φ NMR sample tube by a diaphragm pump, LABOPORT® N86 KN.18 (KNF Neuberger GmbH) at a flow rate of 50 mL/min under atmospheric pressure for HyperCEST NMR/MRI measurements. The polarization level obtained was ~ 10 % using the AW-SEOP-laser system (Aurea Works Corporation) 29. HyperCEST NMR measurements were performed using a pulse sequence: b_1_ - d_1_ - b_2_ - d_2_ - TS - α - tacq where b1 and b2 are gas bubble periods of 1.0 s, d_1_ and d_2_ are delay periods of 0.5 s to make the GV cell suspension static, TS is a radio-frequency (RF) saturation period of 2.0 s, and α is the flip angle of the RF pulse used for observation of dissolved HPXe ^NMR signal (194 ppm) after saturation, t_acq_ is the NMR acquisition period. During the TS^ period, a hundred frequency-selective saturation RF pulses (inter-pulse delay 16 ms) were irradiated. Frequency-dependent saturation spectra were acquired by varying the saturation pulse offset from −83.7 ppm to 278.0 ppm, yielding Z-spectra (i.e. dissolved HPXe signal acquired with off-resonance saturation normalized by that acquired with on-resonance saturation). Z-spectra were acquired from at least four different cell preparations of each GV cell type to ensure reproducibility. Prior to HyperCEST NMR studies, the duration of the saturation and observation pulses (sinc-shaped 4000 μs and Gauss-shaped 800 μs, respectively) was calibrated to achieve saturation. The strength of the saturation RF pulse was 1.5 μT (30dB). Chemical shifts were referenced to free gaseous HPXe NMR signal (0 ppm). Imaging was performed using a four-shot balanced steady-state free precession (b-SSFP) sequence (inter-shot delay 3s) modified to acquire HyperCEST MR images. Frequency saturation similar to the HyperCEST NMR measurement was conducted for 2 s within the inter-shot delay. The duration and strength of the saturation RF pulse employed for HyperCEST MRI were Gauss-shaped 1000 μs and 5.9μT (52dB), respectively. MRI acquisition parameters were as follows: a 1000 μs Gaussian-shaped RF pulse of flip angle α=40º with a bandwidth of 2800 Hz centered on the dissolved HPXe resonance; TR/TE = 3.2ms/1.6ms, acquisition bandwidth, 62 kHz; transverse slice thickness, 20 mm; matrix, 64 × 32 (reconstructed to 128 × 64) with a field of view of 80 × 25 mm^2^; echo train length, 8; number of averages, 16; centrically-ordered phase encoding. HyperCEST NMR/MRI cell samples were prepared in a 15 cm tissue culture dish (IWAKI), peeled with 0.25 % Trypsin-EDTA (Invitrogen), washed with PBS twice and resuspended in PBS with 1/2000 amount of PE-L antifoam agent for cell cultures (FUJIFILM Wako Pure Chemical Corporation) and prepared in custom glass sample tubes.

## Supplementary Information

### Supplementary Text

#### Selection of GV genes

We chose the *gvpC28* gene from *planktothrix agardhii* instead of *planktothrix rubescens* in this study because the full sequence of the *gvpC28* gene was not available on the Genbank database. In addition, because previous studies indicated that a heterologous combination of GV genes could be utilized to induce GVs with altered size and shape into bacterial species ^1^, we assumed that the heterologous combination of GV genes derived from *planktothrix rubescens* and *planktothrix agardhii* could be used to produce GVLPs for these closely related strains.

#### Notes on TEM observation of GV structures

In an initial effort to confirm GVLP expression in the cells, we observed ultra-thin slices of embedded GV_AC20 cell samples by TEM and found the putative GVLPs (indicated white arrow in Supplementary Figure 4), exhibiting the characteristic bicone structure, although the complexity of other intracellular components made it difficult to undeniably confirm GV expression.

It is noteworthy that fixation of the GVLP purified from GV_AC28 cells with 2.5 % glutaralaldehyde (GA) prior to drying and staining was found to be essential for observation of distinct GV structures; without GA fixation, we could not observe distinct GV structures in our experiments (Supplementary Figure 5).

#### Purified GVLP structure and size distribution

The TEM image of GVLPs purified from GV_AC20 cells in Figure 2b shows that the width and length of GVLPs was heterogeneous and ranged from ~ 200 nm to 300 nm. GVLP shape appeared to be biconical or cylindrical without conical ends, comprising a linear outline. The observed cylinders without conical ends may indicate deficient structure formation (e.g. lack of closed protein shell structure); this would reasonably explain the discrepancy between observed TEM sizes (Figure 2b) and lower size distribution peak of about 90 nm in DLS data (Supplementary Figure 2). In contrast, TEM images of GVLPs purified from GV_AC28 cells indicated a more rounded outline compared to other GV cells. The GV size ranged from 100 nm to 700 nm, and included the largest GVs among all that were generated. The two peaks of GVLPs purified from GV_AC28 cells in Dynamic Light Scattering (DLS) data (Supplementary Figure 2) may corresponded to the relatively broad particle size distribution of GVLP in GV_AC28 cells and/or aggregation of the particles. The TEM data showed that the length of GVLPs purified from GV_AC16C20 cells was around 100 nm or less, with a width of 50 - 60 nm. DLS data of GVLPs purified from GV_AC16C20 cells indicated a relatively homogeneous size distribution with a peak at ~ 80 nm, which is in accordance with the TEM data.

### Supplementary Figures

**Supplementary Figure 1.**
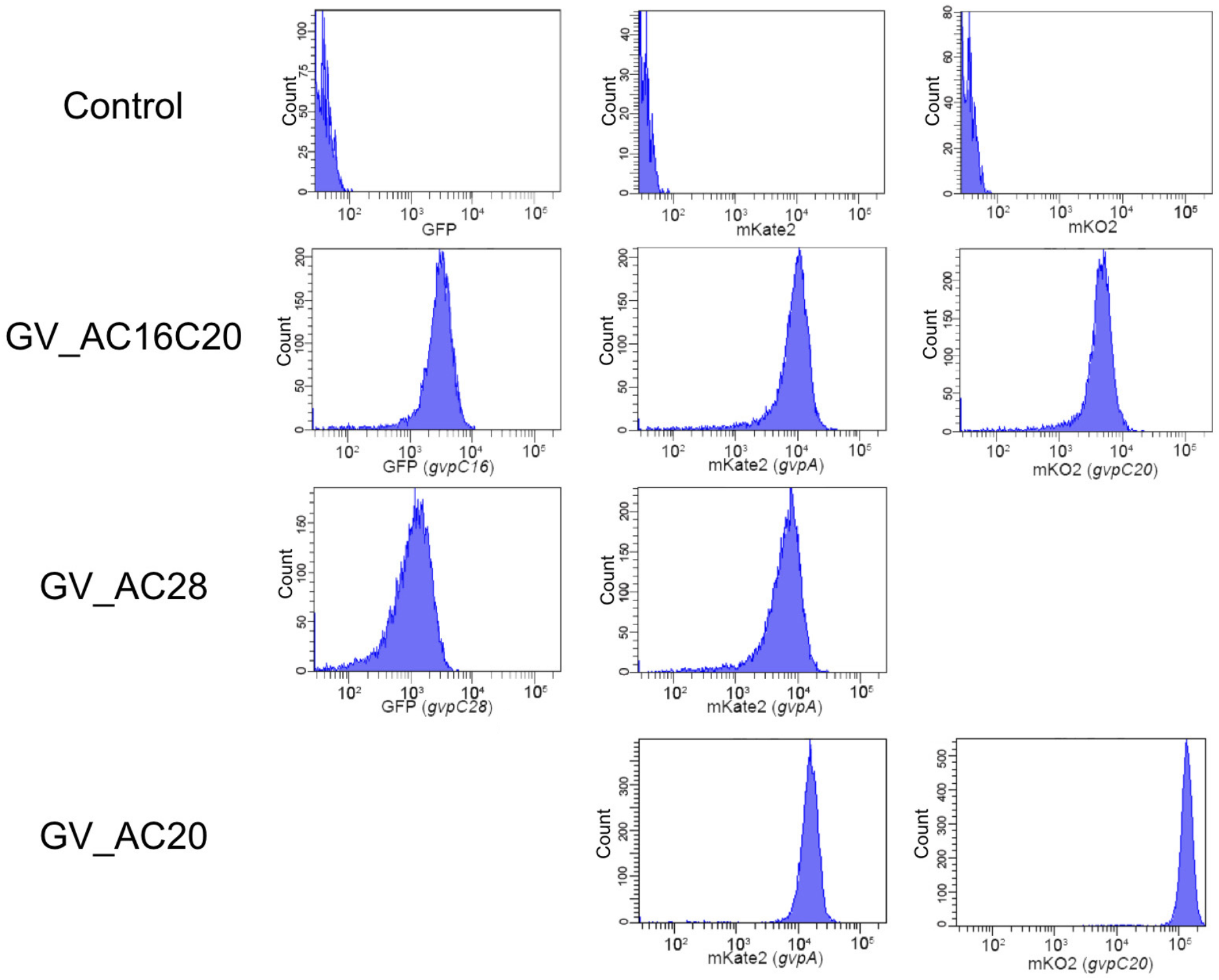
Cell population distribution against fluorescence level, analyzed using FACS and shown as histograms for each GV cell type. The first, second and third column of histograms shows the fluorescence levels of GFP, mKate2 and mKO2, respectively.

**Supplementary Figure 2.**
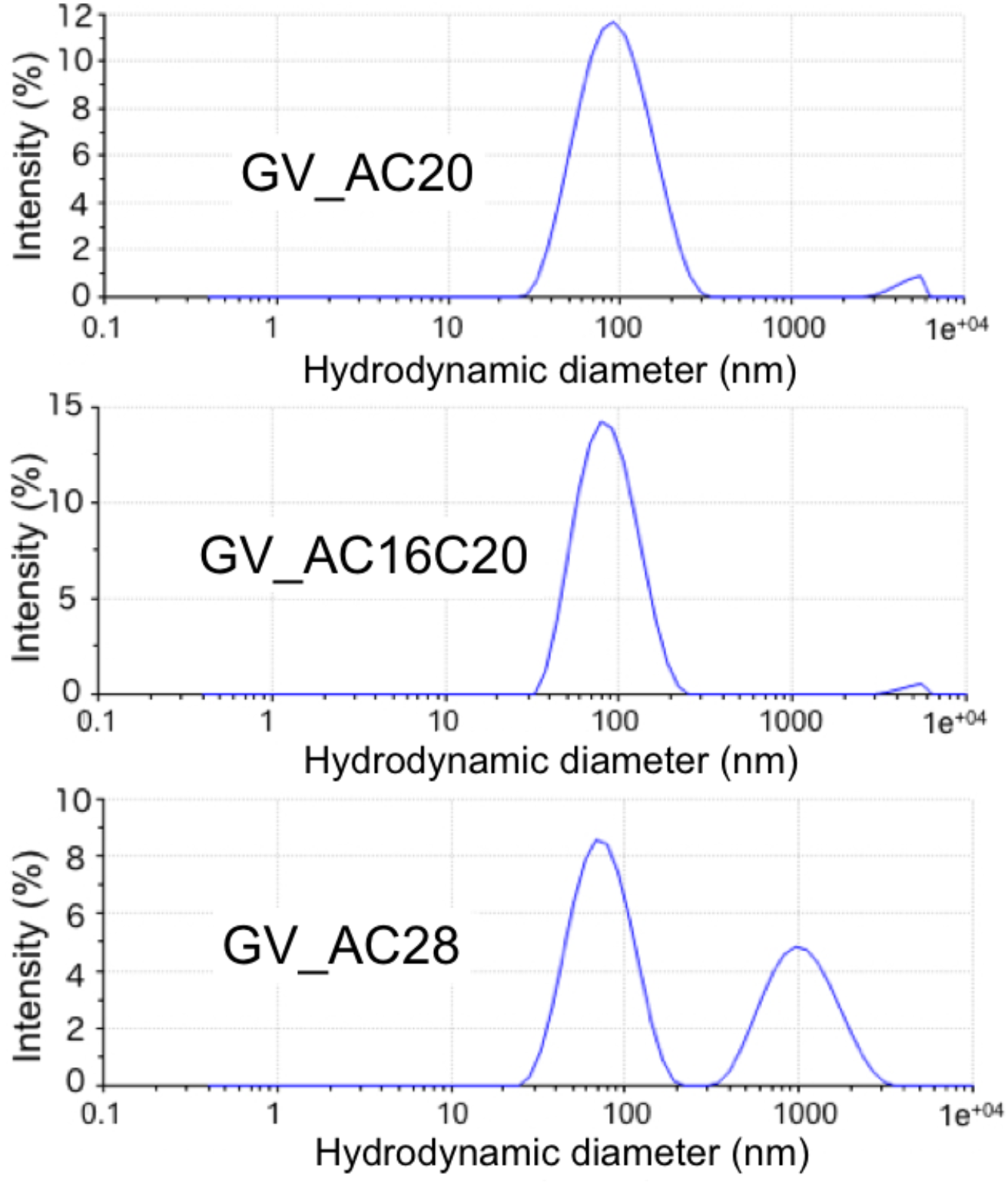
Dynamic Light Scattering (DLS) data on the size distribution of each purified GVLP type. Note that indicated sizes should be interpreted with caution, because GVLPs are non-spherical.

**Supplementary Figure 3.**
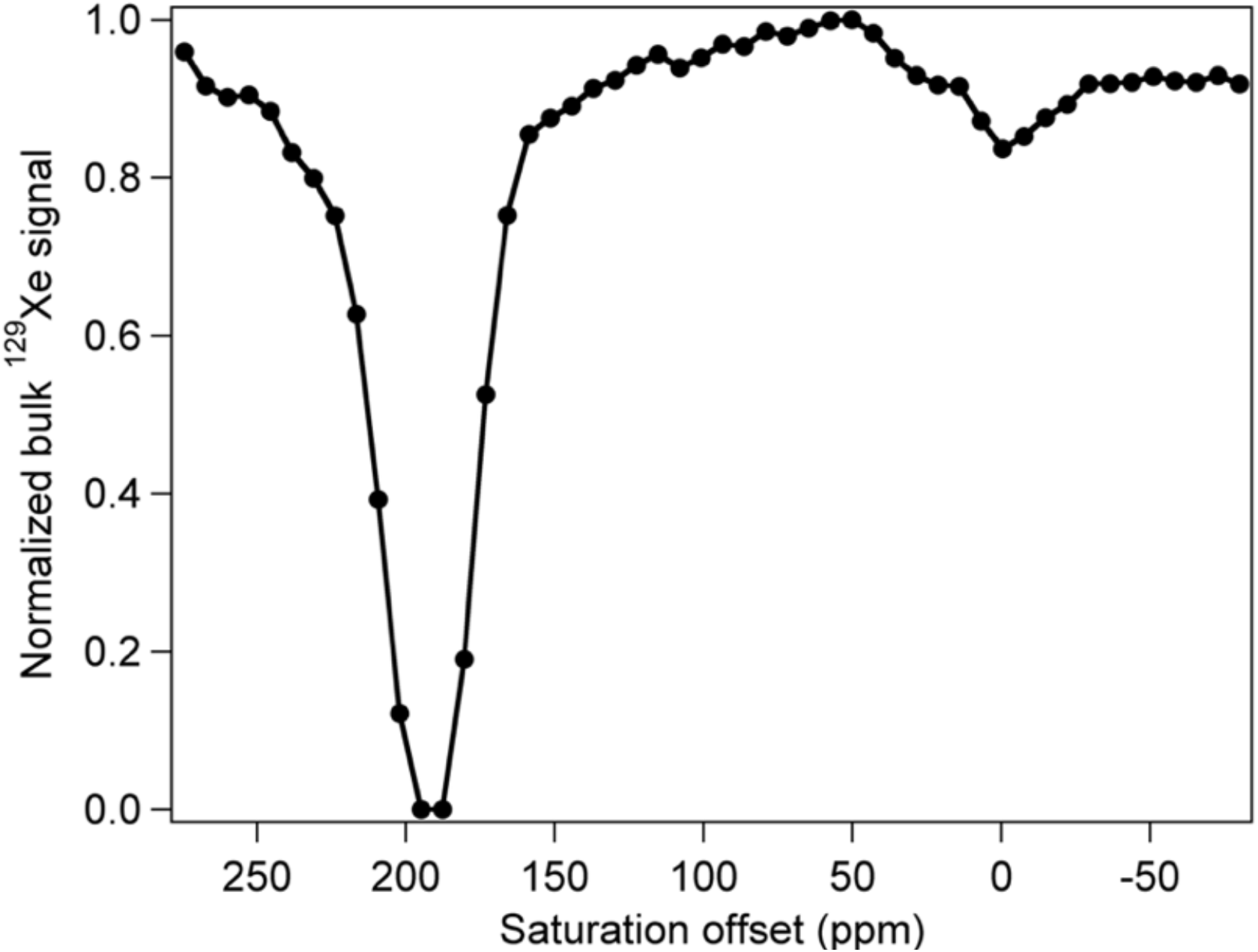
Z-spectrum of a GV_AC20 cell sample at 2 s saturation time with 8.0 × 10^7^ cells / ml. (c.f. Figure 3 of main manuscript)

**Supplementary Figure 4.**
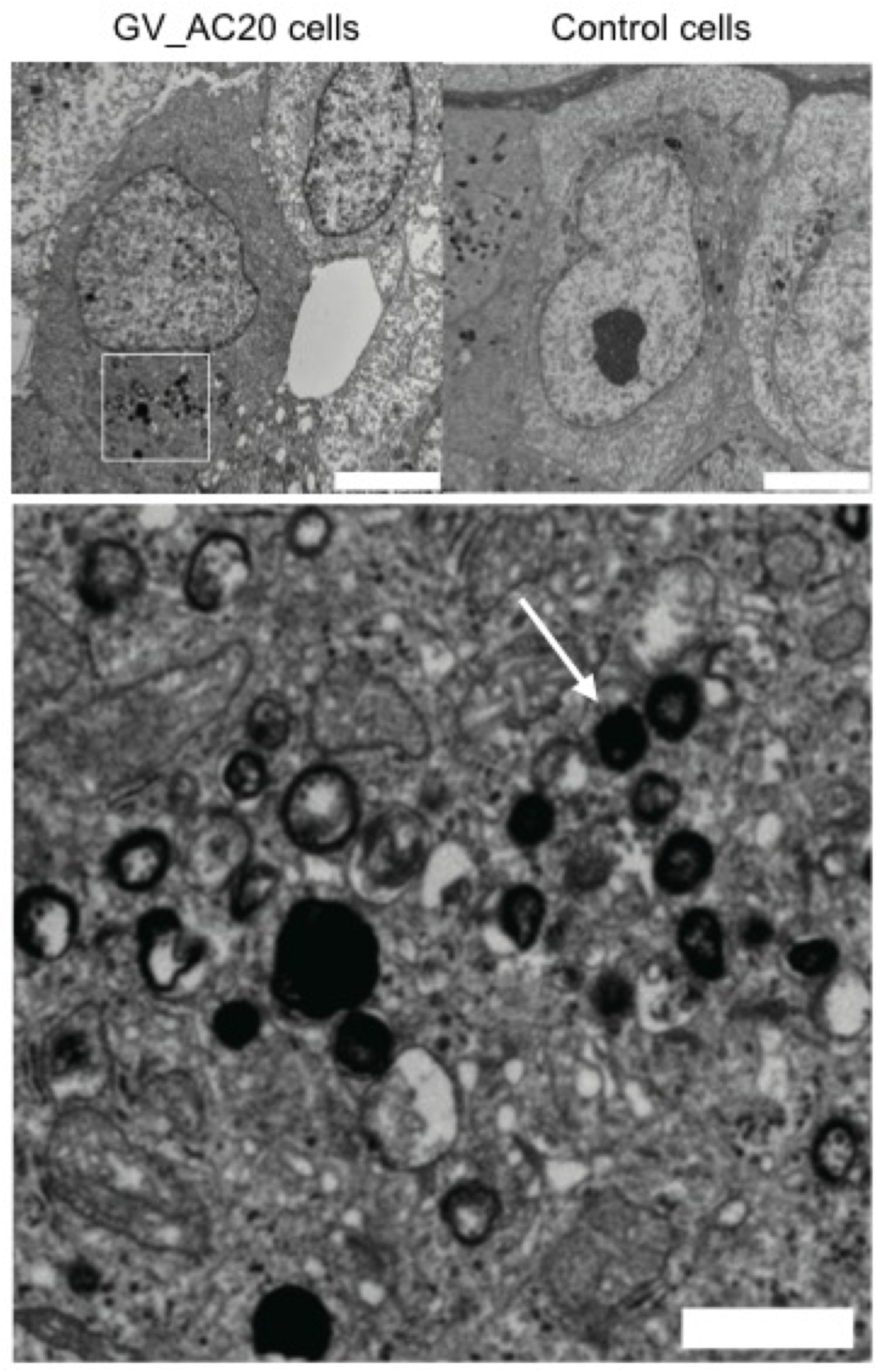
TEM observation of ultra-thin sections of GV_AC20 cells. Top: Sectional TEM images of embedded GV_AC20 (upper left) and control KPL-4 (upper right) cells are shown. Scale bars indicate 5 μm in both images. Bottom: Magnified region of interest (depicted by white square in upper left image) of the GV_AC20 cell image. The scale bar indicates 1 μm. The white arrow indicates a biconical gas vesicle.

**Supplementary Figure 5.**
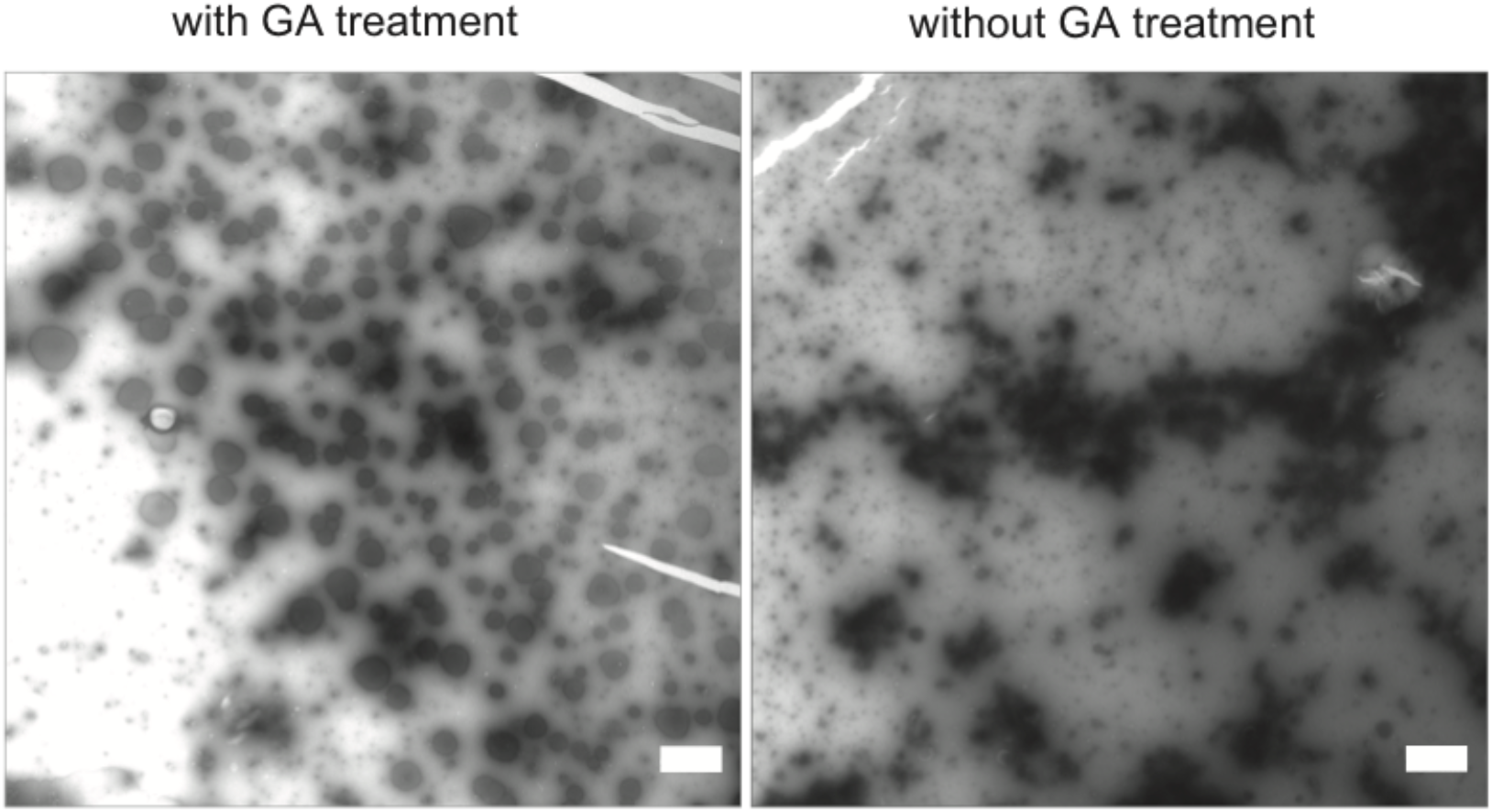
TEM images of GVLPs purified from GV_AC28 cells with and without glutaraldehyde (GA) treatment prior to negative staining. Scale bars indicate 1 μm in both images. These images illustrate the importance of the GA treatment for visualization of GV structures.

## Notes

https://dnaconda.riken.jp/search/depositor/dep103337.html

